# The Impact of Ketamine and Thiopental Anesthesia on Ultraweak Photon Emission and Oxidative-Nitrosative Stress in Rat Brains

**DOI:** 10.1101/2024.08.22.609284

**Authors:** Mahdi Khorsand Ghaffari, Niloofar Sefati, Tahereh Esmaeilpoor, Vahid Salari, Daniel Oblak, Christoph Simon

## Abstract

Anesthetics such as ketamine and thiopental, commonly used for inducing unconsciousness, have distinct effects on neuronal activity, metabolism, and cardiovascular and respiratory systems. Ketamine increases heart rate and blood pressure while preserving respiratory function, whereas thiopental decreases both and can cause respiratory depression. This study investigates the impact of ketamine (100 mg/kg) and thiopental (45 mg/kg) on ultraweak photon emission (UPE), oxidative-nitrosative stress, and antioxidant capacity in isolated rat brains. To our knowledge, no previous study has investigated and compared UPE in the presence and absence of anesthesia. Here, we compare the effects of ketamine and thiopental anesthetics with each other and with a non-anesthetized control group. Ketamine increased UPE, lipid peroxidation, and antioxidant enzyme activity while reducing thiol levels. Conversely, thiopental decreased UPE, oxidative markers, and antioxidant enzyme activity, while increasing thiol levels. UPE was negatively correlated with thiol levels and positively correlated with oxidative stress markers. These findings suggest that the contrasting effects of ketamine and thiopental on UPE are linked to their differing impacts on brain oxidative stress and antioxidant capacity. This research suggests a potential method to monitor brain oxidative stress via UPE during anesthesia, and opens up new ways for understanding and managing anesthetic effects.

**Graphical Abstract:** 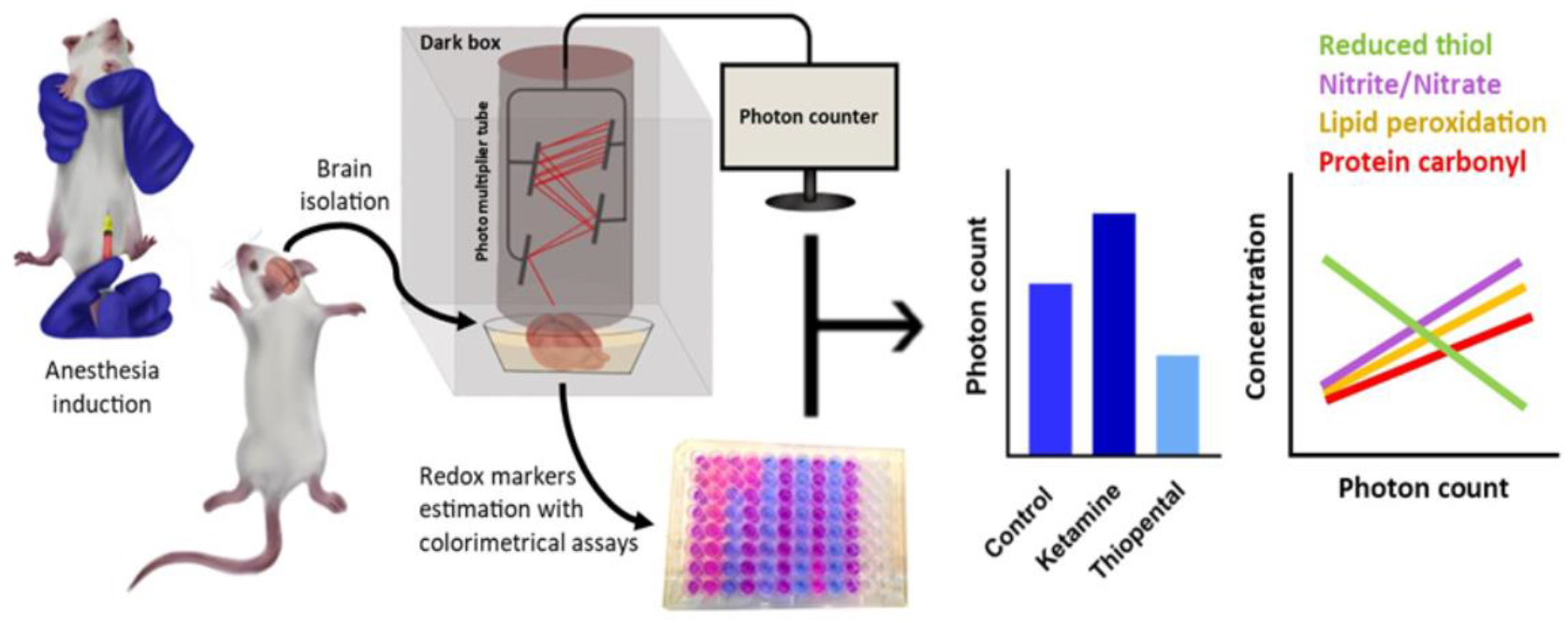

## 1 Introduction

Anesthetics induce regulated unconsciousness associated with extensive brain network connectivity perturbations. They are used for analgesia, sedation, and hypnosis during surgery and in the intensive care unit. The mechanism of anesthesia is still not fully understood. This may be due to the ilusive nature of consciousness (1). Several neuroscientific theories of consciousness have been proposed and are currently the subject of heated debate. Although these ideas may seem different at first glance, they are all fundamentally rooted in various aspects of interconnectedness. Their shared belief is that the richness of neuronal interactions in corticothalamic systems is crucial for achieving consciousness. Specifically, when interacting neurons reach a critical level of complexity, a conscious experience is formed (2).

Large-scale neural oscillations are intricately linked to essential cognitive functions, including perception, attention, decision-making, memory, and consciousness (3). In a study, it has been shown that different spectral light stimulations at one end of the spinal sensory or motor nerve roots significantly increased the biophotonic activity at the other end (4). Thus, local anesthetic or metabolic inhibitors may inhibit these effects. Ultraweak photon emission (UPE) is closely linked to reactive oxygen species (ROS), with intensity variations reflecting various physiological and pathological states, such as stress (thermal, chemical, mechanical), mitochondrial function, cell cycle, etc., and UPE intensity correlates with neural activity, oxidative reactions, EEG activity, cerebral blood flow, energy metabolism, and glutamate release (5). Different anesthetics produce diverse EEG signatures linked to their molecular targets and affected neural circuits. For example, Thiopental dose-dependently decreases EEG frequency until the EEG becomes isoelectric, while ketamine increases high-frequency gamma waves (6, 7). High-frequency oscillations are linked to neuronal firing and cortical activation, while low-frequency oscillations occur during the resting state (8). Therefore, anesthetics and their effect on UPE might be somehow correlated with brain function, EEG, and cognitive behaviours (9, 10).

Neural communication depends on the release of neurotransmitters. This intricate process involves the transportation of molecules to presynaptic active zones by motor proteins, regulation of ion channel function, mobilization of synaptic vesicles, and recycling of neurotransmitters. Each of these steps requires energy in the form of ATP, and general anesthetics can impact these essential processes (11-13). The predominant energy expenditure in the brain involves maintaining ion gradients across cellular membranes, which is largely fuelled by the oxidative metabolism of glucose. The energy demands of glutamatergic neurons account for 80-90% of cortical glucose consumption (14). Astrocytes primarily remove released glutamate from the synaptic cleft. A study has shown a close coupling between cerebral glucose metabolism and the cycling of glutamate between neurons and astrocytes (15).

Anesthesia induction with ketamine and thiopental has different effects on glutamatergic transmission. Ketamine inhibits NMDA receptors of cortical interneurons more efficiently than those on pyramidal neurons. This inhibition leads to reduced interneuron activity and subsequently decreases the release of GABA on pyramidal neurons. The reduction in GABA release ultimately results in increased glutamate release and heightened excitation of pyramidal neurons. Additionally, ketamine may interrupt the steady inhibition of glutamate release by inhibiting presynaptic NMDA receptors (16, 17). Thiopental is a GABA_A_ receptor agonist which depresses cerebral activity by prolonging the opening of the Cl-channel and hyperpolarization induction. Moreover, it has been shown that thiopental sodium inhibits glutamate release from cerebrocortical slices (18, 19). These aforementioned mechanisms explain why ketamine heightens cerebral glucose metabolism while thiopental decreases it (20, 21).

UPE has been detected during the cellular metabolism of all living systems. UPE arises from the electronic relaxation of excited species formed during reactions of reactive oxygen species (ROS) and reactive nitrogen species (RNS) and their derivatives (22). The interaction of biomolecules (lipids, proteins, nucleic acids) with these reactive species creates high-energy intermediate products that eventually lead to the formation of singlet or triplet excited state carbonyl compounds and singlet ground state oxygen. When these species undergo electronic transitions, photons are emitted at short and long wavelength regions of the spectrum, respectively (23).

ROS and RNS primarily originate from the respiratory chain of mitochondria, NADPH oxidases, nitric oxide synthases, xanthine oxidase, cytochrome P450 oxidases, lipoxygenases, and cyclooxygenases. Respiration generates oxygen-centered free radicals, which are molecules with unpaired electrons. The reduction of molecular oxygen to water involves the addition of four electrons in several steps. Molecular oxygen (O_2_) is first reduced to form the superoxide anion radical (O_2_-), which then reacts with its conjugate acid to produce hydrogen peroxide (H_2_O_2_). Hydrogen peroxide is a two-electron reduction product of O_2_ and serves as both a byproduct and a source of free radical reactions. It can also react with iron or other transition metals to produce the hydroxyl radical (OH), a powerful oxidant. Additionally, Superoxide can react with nitric oxide (NO) to form the peroxynitrite anion (OONO−). When protonated to peroxynitrous acid (OONOH), it decomposes to yield the hydroxyl radical (24). Superoxide dismutase (SOD) facilitates the conversion of O_2_-into H_2_O_2_. The catalase enzyme (CAT) subsequently breaks down H_2_O_2_ into H_2_O and O_2_, or it can be transformed by thiol peroxidases, such as glutathione peroxidase, which promote the reduction of H_2_O_2_ and/or organic hydroperoxides into water and the corresponding alcohols. Unlike SOD and CAT, peroxidases utilize a thiol-based reaction mechanism to neutralize hydroperoxides. Thiol reduced state must be replenished by reductases, such as glutathione reductase, with the consumption of NADPH (25).

Thiol is an organosulfur compound in the form R−SH, where R represents an alkyl or other organic substituent, and −SH is a functional sulfhydryl group. Cysteine amino acid represents this characteristic. Organisms create millimolar concentrations of cysteine-containing thiols that serve as a cofactor for thiol-dependent enzymes. Moreover, they form covalent linkages with protein thiols to protect against overoxidation and reduce existing oxidative modifications in proteins. The tripeptide γ-glutamyl cysteinyl glycine, known as glutathione (GSH), appears to maintain cellular redox homeostasis in most eukaryotes (26). When ROS are effectively scavenged by GSH, their oxidative effects on biomolecules are prevented. However, biomolecules are oxidized if ROS formation exceeds the antioxidant defence system’s capacity. Studies have shown that the addition of exogenous GSH, SOD, and CAT suppressed ultra-weak photon emission (27, 28).

Ketamine and thiopental anesthetics have the opposite effect on brain metabolism when assessed using neuroimaging techniques. However, these methods may be affected by respiratory and cardiovascular interference (29). Studies have shown that ketamine and thiopental have opposite respiratory and cardiovascular effects (30). Isolated brain involves submerging the brain in oxygenated artificial cerebrospinal fluid. This approach maintains brain integrity without perfusing the vascular system. The viability of the rat brain using this method was confirmed by the successful recording of extracellular field potentials one day later (31). Moreover, the EEG activity of the isolated brain was recorded and could last for up to 30 minutes (32).

To our knowledge, no previous study has investigated and compared UPE in the presence and absence of anesthesia. Given that UPE is closely linked to oxidative metabolism and considering that anesthesia can alter brain metabolism and activity, we examine the isolated brain’s UPE under anesthesia with ketamine and thiopental. We also evaluate the relationship between changes in UPE and the brain’s oxidative-nitrosative state and antioxidant capacity.

## 2 Material and methods

### 2.1 Drugs and reagents

Ketamine (Vetased^R^, Pasteur, Romania), Thiopental, Ethylenediaminetetraacetic acid (EDTA), thiobarbituric acid (TBA), 1,1,3,3-tetraethoxypropane (TEP), vanadium (III) chloride (VCL3), sulfanilamide, N-(1-Naphthyl) ethylenediamine dihydrochloride (NEDD), sodium nitrite (NaNO2), 2,4-dinitrophenylhydrazine (DNPH), trichloroacetic acid (TCA), guanidine hydrochloride, 5,5-Dithiobis-2-nitrobenzoic acid (DNTB). Except for ketamine, all other chemicals were purchased from Sigma-Aldrich.

### 2.2 Artificial cerebrospinal fluid (aCSF) composition

The aCSF contains 124mM NaCl, 3mM KCl, 26 mM NaHCO3, 1.25 mM Na2HPO4, 1.8 mM MgSO4, 1.6 mM CaCl2, 10 mM glucose

### 2.3 Animals

Eighteen male Sprague-Dawley (SD) rats weighing between 180-190 g were randomly assigned to three experimental groups: Control, Ketamine, and Thiopental (n=6). Rats were purchased from the Comparative and Experimental Medical Center of the Shiraz University of Medical Sciences (SUMS). All rats were kept in standard conditions, including a temperature of 22±2 °C, relative humidity of 50%, and a 12-hour light/dark cycle, with free access to laboratory food and water. Animal procedures complied with the National Institutes of Health’s Guide for the Care and Use of Laboratory Animals and the ARRIVE Guidelines. The university’s Ethics Committee approved the procedures (approval number: IR.SUMS.REC.1400.191).

### 2.4 Study design

The control group received an intraperitoneal (IP) injection of 5 ml/kg of saline. After 3 minutes, the head was dislocated, and the brain was quickly removed to minimize any oxidative and anesthetic interventions. Other rats were given IP ketamine or thiopental. Following injection, anesthesia induction was checked with the loss of three successive righting reflexes. Then, their heads were dislocated, and the brain was rapidly removed. A study found that Sprague-Dawley rats had similar anesthesia induction and maintenance times with IP injection of 100mg/kg ketamine or 45mg/kg thiopental (33). Therefore, these doses were selected for comparison in the present study. After the brain was removed, it was flooded into a chamber containing fresh oxygenated aCSF (95% O2, 5% CO2). To prevent any possible delayed luminescence, the chamber was left in a dark room for 10 minutes while being continuously oxygenated with carbogen at room temperature (34). After that, the brain was transferred to a chamber filled with fresh oxygenated aCSF at room temperature (∼30°C) and positioned under a photomultiplier tube (PMT) to detect UPE for 300 seconds. Next, the brain was fast-frozen with liquid nitrogen and stored at -80 for further assessment of reduced thiol, lipid peroxidation, nitrite/nitrate levels, protein carbonyl, and antioxidant enzyme activity (SOD, CAT).

### 2.5 UPE detection

In this study, a PMT (R6095 Hamamatsu Photonics K.K., Japan) was used to detect UPE in a dark box. The PMT has a quantum efficiency (QE) that varies between 20 and 30% in the range 300-700 nm, with the maximum QE at 420 nm. PMT amplifies entrance photons to electrical signals in a field of view. The analog signals were converted to digital using an RS485 to RS232 converter and then connected to a laptop for recording. The collecting gate time from the PMT was set to 1 second. Noise is reduced by modifying the upper and lower thresholds via PMT software. Dark noise was detected in an empty box for 5 minutes (∼ 1100 counts per 5 min) and subtracted from the results. The distance between the sample and the PMT sensor was adjusted to 0.5 cm. Before each trial, the aCSF emissions were recorded for 5 minutes and subtracted from sample emissions.

### 2.6 Redox markers assessment

#### 2.6.1 Brain homogenate preparation

The frozen brain was weighed and then homogenized using a Homogenizer (T 10 basic ULTRA-TURRAX, IKA, Germany) in ice-cold EDTA-potassium phosphate buffer for around 3 minutes. After homogenization, the mixture was centrifuged at 12000 rpm at 4°C for 5 minutes. The resulting supernatant was then separated and stored at -80°C for further colorimetric measurements with a microplate reader (BioTek Synergy H1, Agilent, USA).

#### 2.6.2 Estimation of lipid peroxidation level

The level of lipid peroxidation in animal tissues was measured using the TBA reaction method. In this method, the pink color produced during the reaction of TBA with peroxidized lipids was measured at 532 nm for the estimation of lipid peroxidation, and the TEP was used as a standard (35). TBA stands for “thiobarbituric acid”, a chemical used in the “thiobarbituric acid reactive substances (TBARS)” assay to measure lipid peroxidation. The TBA reacts with malondialdehyde (MDA), a byproduct of lipid peroxidation, forming a pink chromogen that can be quantified by measuring absorbance at 532 nm. TEP stands for “1,1,3,3-Tetraethoxypropane”, which is commonly used as a standard in lipid peroxidation assays like the TBARS assay. It is used because TEP can generate MDA when hydrolyzed, which then reacts with TBA to form the pink chromogen measured at 532 nm. By comparing the absorbance of sample to that of known TEP standards, one can quantify the amount of lipid peroxidation in your samples.

#### 2.6.3 Estimation of nitrite/nitrate level

The nitrite/nitrate level is estimated based on the previously described colorimetric method (36). First, nitrate is converted to nitrite with a reaction to VCL3. Subsequently, the nitrite reacts with sulfanilamide at low pH to produce a diazonium salt. This salt is then reacted with NEDD to form a stable compound. The nitrite/nitrate level of the sample is estimated by comparing the absorbance of this compound with a standard curve of NaNO2 at 540 nm.

#### 2.6.4 Estimation of Protein carbonyl

The protein’s oxidative damage was assessed by measuring the presence of carbonyl groups, which could be estimated by reacting them with DNPH. As described by Levine (37), DNPH in hydrochloride was added to the homogenate supernatant and incubated for 1 hour. Subsequently, proteins were precipitated by adding TCA. After centrifugation, the supernatant was removed, and the pellets were washed with ethanol and ethyl acetate to eliminate excess DNPH. They were then dissolved in a solution of guanidine hydrochloride, and the absorbance was measured at 375 nm. Protein carbonyl level was determined using a molar extinction coefficient of 22,000 M−1 cm−1.

#### 2.6.5 Estimation of reduced thiol concentration

The concentration of reduced thiol was determined by a colorimetric method. In this method, Ellman’s reagent, also known as DNTB, reacts with reduced sulfhydryl groups. The complex formed is called the 5-thionitrobenzoic acid chromophore. This complex produces a yellow color, and the absorbance at 405 nm is used to estimate the concentration of reduced thiol in the sample (38).

#### 2.6.6 Superoxide dismutase and Catalase activity estimation

SOD and CAT activity were measured using the colorimetric method, according to the manufacturer’s instructions of the Assay Kit (ZellBio GmbH, Germany).

#### 2.6.7 Supernatant protein concentration

The Bradford method was used to assess the total protein content in the tissue supernatant (39). Molecular data were expressed as x/mg protein of the supernatant to normalize the concentration.

### 2.7 Statistical analysis

Data were expressed as the means ± standard error of the mean (SEM). Statistical significance was set at P < 0.05. A one-way ANOVA followed by Tukey’s post hoc test was used to compare the results among different groups. The relationship between UPE and oxidative-nitrosative stress was evaluated by Pearson correlation analysis. All statistical analyses were conducted using GraphPad version 8 (Prism Software Inc., San Diego, CA, USA).

## 3 Results

### 3.1 Effects of anesthesia induction on brain UPE

In this study, we found that anesthesia induction significantly altered the UPE of the isolated brain. Specifically, ketamine increased UPE while thiopental decreased it when compared to the control group (p < 0.05; Fig 1). In comparison between anesthesia groups, the thiopental group had lower brain UPE than the Ketamine group (p < 0.0001).

**Fig 1:**
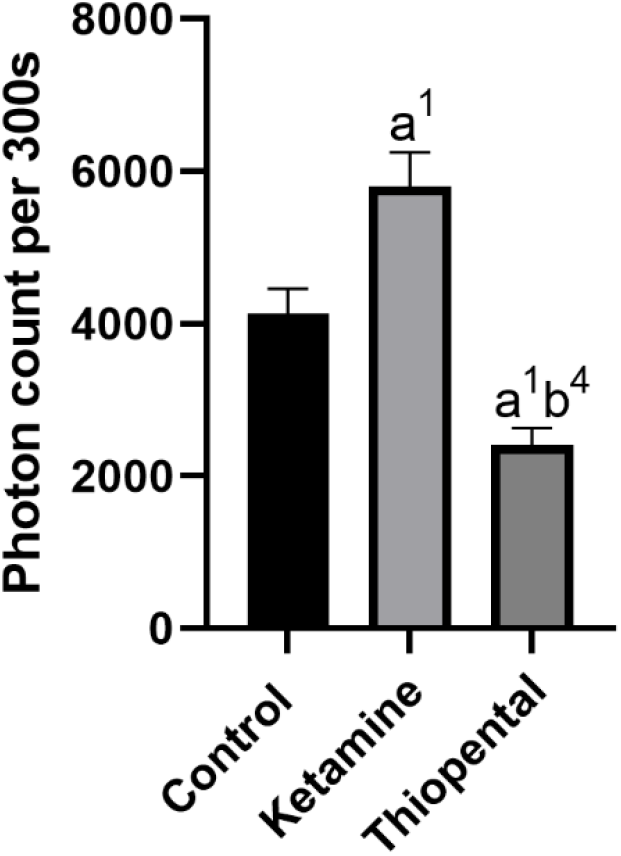
Number of brain’s UPE counted in the 300 seconds. Data were presented as mean ± SEM (n=6) and analyzed by one-way ANOVA followed by Tukey’s multiple comparison test. (a) compared to the control group, and (b) compared to the ketamine group. 1p < 0.05, 4p < 0.0001.

### 3.2 Effects of anesthesia induction on brain oxidative and nitrosative state

Results from the oxidative and nitrosative state of the isolated brain showed that ketamine significantly increased lipid peroxidation (Fig 2A) and nitrite/nitrate level (Fig 2C) as compared to the control group (p < 0.05, p < 0.0001, respectively). In contrast, thiopental decreased the lipid (p < 0.01; Fig 2A), protein (p < 0.05; Fig 2B) and nitrogen (p < 0.01; Fig 2C) oxidation of the brain as compared to the control. In comparison between anesthesia groups, the thiopental group had a significantly lower brain oxidation state than the ketamine group (p < 0.01; Fig 2).

**Fig 2:**
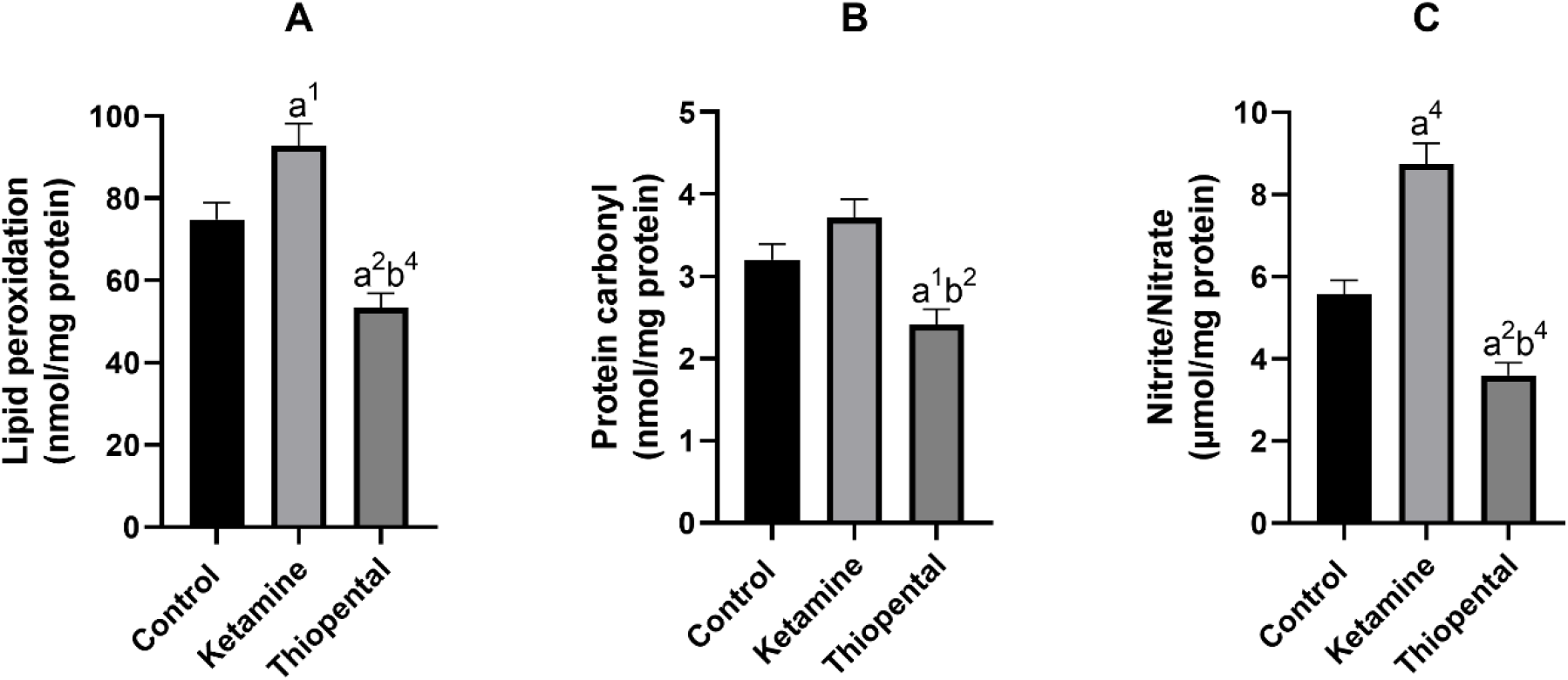
Brain oxidative and nitrosative state. (A) Lipid peroxidation level of the brain, (B) protein carbonyl content of the brain, (C) nitrite/nitrate level of the brain. Data were presented as mean ± SEM (n=6) and analyzed by one-way ANOVA followed by Tukey’s multiple comparison test. (a) compared to the control group, and (b) compared to the ketamine group. 1p < 0.05, 2p < 0.01, 4p < 0.0001.

### 3.3 Changes in the brain’s antioxidant capacity under anesthesia

As shown in Fig 3A, the brain reduced thiol level of the ketamine group was significantly lower than the control (p < 0.001). Brain antioxidant enzyme activity assessment indicated that anesthesia induction with ketamine increased brain SOD (Fig 3B) and CAT (Fig 3C) activity as compared to the control (p < 0.05). In contrast, anesthesia induction with thiopental increased reduced thiol level and decreased SOD activity compared to the control (p < 0.01, p < 0.05, respectively). In comparison between anesthesia groups, the thiopental group had higher reduced thiol (p < 0.0001; Fig 3A) and lower SOD (p < 0.0001; Fig 3B) and CAT activity (p < 0.01; Fig 3C) than the ketamine group.

**Fig 3:**
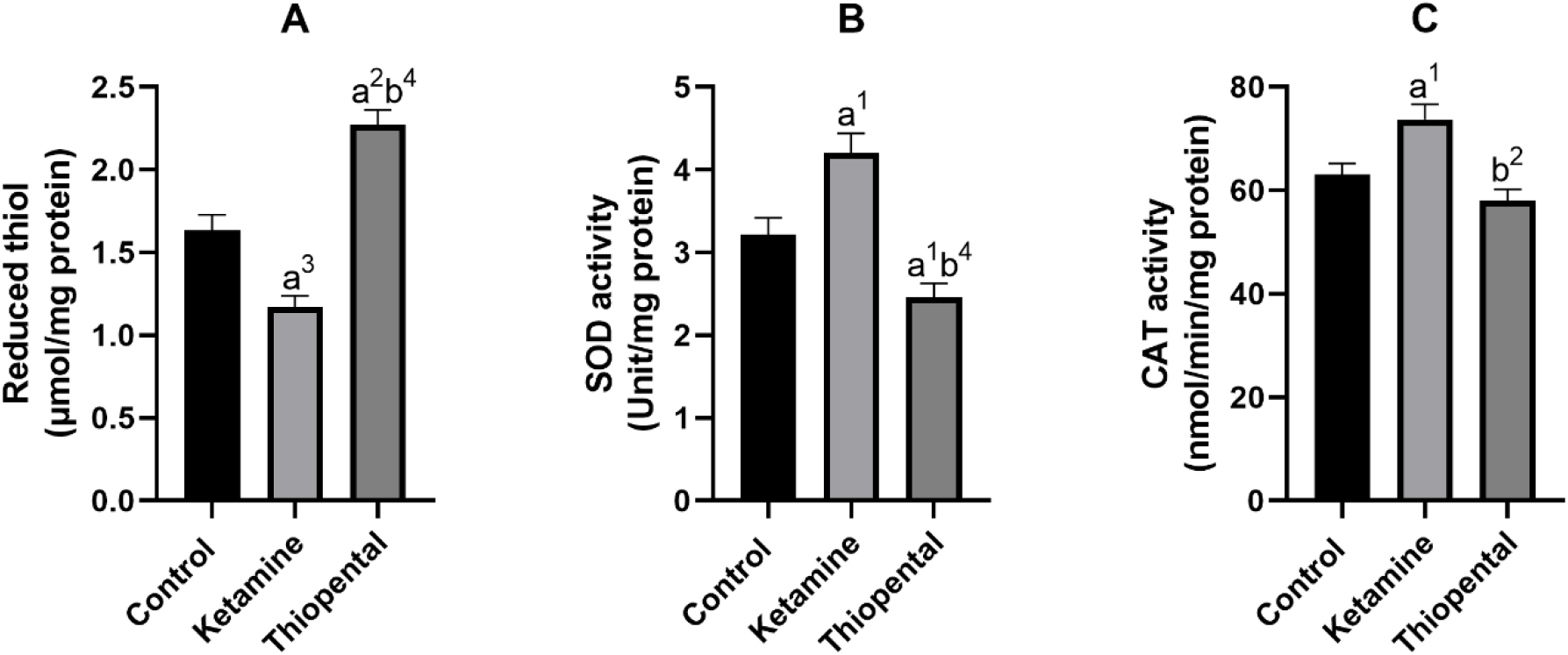
Antioxidant capacity of the brain. (A) reduced thiol level of the brain, (B) SOD activity of the brain, (C) CAT activity of the brain. Data were presented as mean ± SEM (n=6) and analyzed by one-way ANOVA followed by Tukey’s multiple comparison test. (a) compared to the control group, and (b) compared to the ketamine group. 1p < 0.05, 2p < 0.01, 3p < 0.001, 4p < 0.0001.

## 4 Discussion

The purpose of this study is to evaluate the isolated brain’s UPE (ultraweak photon emission) under anesthesia induced by ketamine and thiopental. The findings indicate that anesthesia induction can indeed alter the UPE of the isolated brain. To further explore the relationship between changes in UPE and the brain’s oxidative-nitrosative stress and antioxidant capacity, we conducted data analysis using Pearson correlation analyses. These analyses showed positive relationships between UPE and lipid (Fig 4A) protein (Fig 4B), and nitrogen (Fig 4C) oxidation of the brain (r = 0.8249, r = 0.7719, r = 0.8586, respectively). Moreover, a negative relation between UPE and the reduced thiol level of the brain was detected (r = -0.8736; Fig 4D). UPE had the strongest correlation with reduced thiol level and the weakest correlation to protein carbonyl. The Coefficient of determination (r2) indicated that 76% of the UPE variance could be explained by reduced thiol, while 59% of UPE variance could be explained by protein carbonyl.

**Fig 4:**
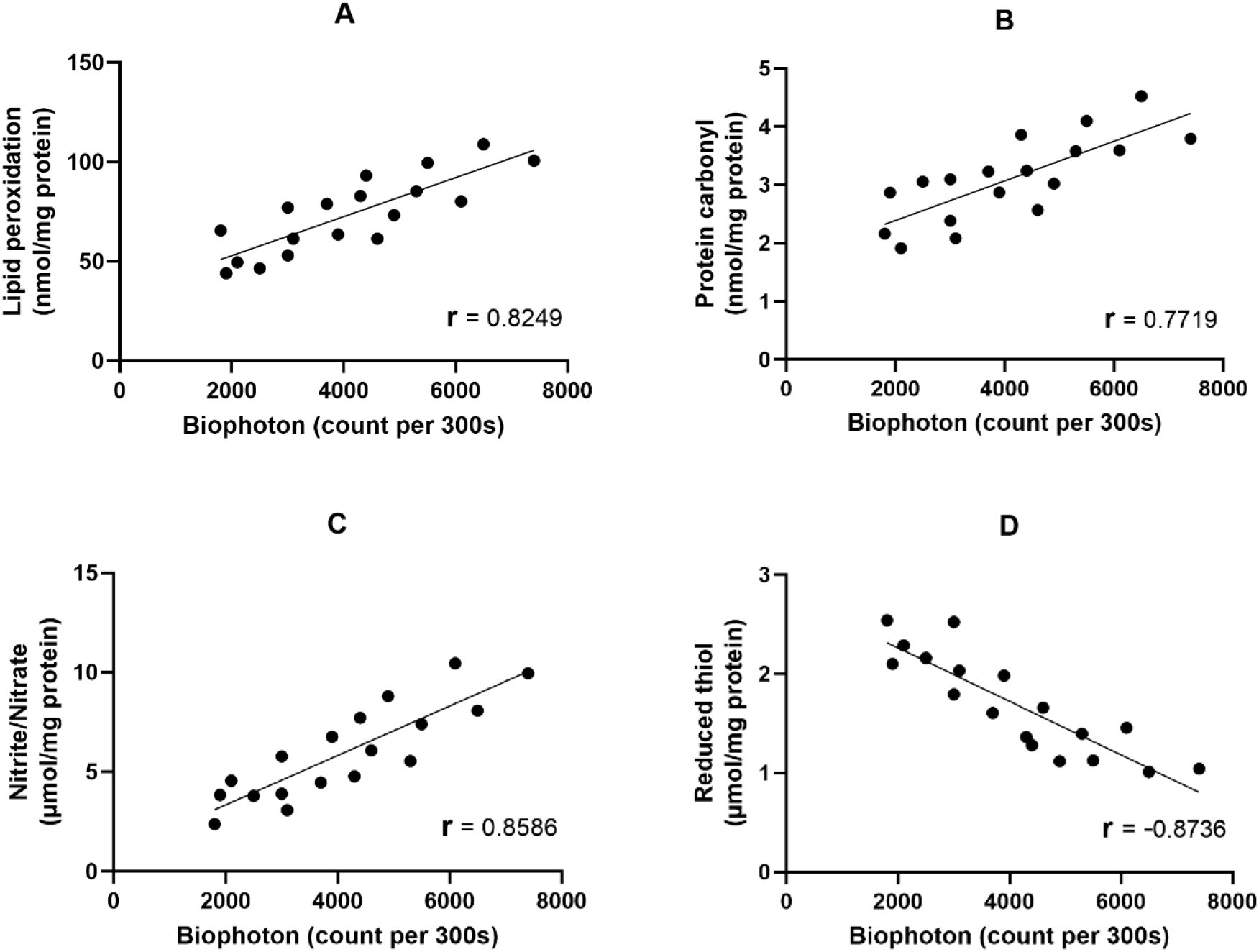
Correlation matrix and linear regression of the isolated brain UPE with oxidative stress markers. (A) UPE vs. Lipid peroxidation; (B) UPE vs. Protein carbonyl; (C) UPE vs. Nitrite/Nitrate; (D) UPE vs. Reduced thiol.

In agreement, it was shown that the intensity of rat brain UPE is correlated to oxidative stress which arises from electrical activity and energy metabolism (40). Studies have demonstrated that ROS and RNS through the oxidation of lipid, protein, and nucleic acid produce excited species. Electronic relaxation of these species appears to be the source of UPE. Mitochondria are thought to be the main source of this activity. Since the respiratory chain of mitochondria contains a large amount of molecular oxygen it is predisposed to radical generation. Therefore, photon emission intensity is closely coupled to oxidative metabolic activity (23, 41).

Neuron firing and communication consume ATP, and, hence, scale-up metabolism. Glutamate is the main excitatory neurotransmitter in the CNS. Tang et al, showed that the glutamate application on brain slices intensified biophotonic activities and this activity could be blocked by oxygen and glucose deprivation (42). Ketamine and thiopental have contrasting effects on glutamate release and brain metabolism. It was shown that ketamine through blocking the NMDA receptors on inhibitory interneurons, leads to excitatory activity and glutamate release from downstream pyramidal neurons (16). A recent study demonstrated that an anesthetic dose of ketamine broadly enhanced brain metabolism in SD rats, which makes sense given that cerebral metabolism and neuronal activity are connected (43). In contrast, it was found that the spontaneous and KCl-evoked release of glutamate in the prefrontal cortical synaptosomes was suppressed by thiopental and it reduced the extracellular level of glutamate in the rat prefrontal cortex (44, 45). ^13^C magnetic resonance spectroscopy of rat cortex under anesthesia with thiopental showed a significant decrease in the oxidative metabolism of neurons and astrocytes (46). As biophotonic activity correlated to glutamate and brain metabolism, consistent with aforementioned evidences, we found ketamine increased the brain UPE while thiopental decreased it (Fig 1).

Ketamine by antagonizing NMDA receptors on interneurons reduces the inhibitory effect of them on monoaminergic neurons. The absence of this inhibitory input leads to an increase in dopamine, serotonin, and norepinephrine release (48-50). On the contrary, it has been shown that thiopental decreases the release of dopamine, serotonin, and norepinephrine (50, 52, 53).

It was proposed that increased UPE production and decreased UPE absorption and transfer may cause cellular oxidative damage (56). Hence, both ROS and UPE could lead to an increase in oxidative damage. ROS is generated by the interaction of superoxide anion with other molecules. This anion is created by electron addition to molecular oxygen in the respiratory chain of mitochondria and Complex I is the major source of this activity (57). Venâncio et al. showed that an anesthetic dose of ketamine disrupted Complex I function, leading to elevated oxygen consumption, reduced efficiency of oxidative phosphorylation, enhanced H_2_O_2_ generation, and nitric oxide synthase by mitochondria. They suggested that this occurred due to reversed electron transport, which is linked to the generation of superoxide and nitric oxide (58). Consistent with this evidence previous studies showed that shortly after a single dose of ketamine, lipid peroxidation, nitrite content, protein carbonyl, SOD activity, and CAT activity of the brain increased, while the reduced thiol level decreased (58-61). In agreement, we found these changes in the ketamine group (Fig 2 and 3).

Increased antioxidative enzyme activity in the ketamine group may result from increased expression of them due to NO-induced extracellular-regulated kinase 1/2 pathway or H2O2-induced translocation of the nuclear factor κB into the nucleus (62, 63). A study found that the expression of stress proteins such as SOD1 increased shortly after exposure to oxidative stress (64).

Protein synthesis was identified as the most ATP-consuming process in mammalian cells (65). Thiopental was demonstrated to inhibit global protein synthesis in neurons by inactivating eukaryotic elongation factor 2. Through this mechanism, it preserves ATP and prevents damage in oxygen-deprived cells (66). This evidence could explain reduced SOD and CAT activity in the thiopental group (Fig 3B, C).

Protein synthesis inhibition by thiopental may also block the translation of inducible nitric oxide synthase, cyclooxygenase-2, or matrix metalloproteinases. These proteins have been implicated in peroxynitrite formation, lipid peroxidation, and protein oxidation (67). In agreement, we found a decrease in lipid peroxidation, protein carbonyl, and nitrite/nitrate levels in the brain of the thiopental group (Fig 2). Previous studies have demonstrated that thiopental exhibits a potent capacity to scavenge all types of radicals and could prevent lipid peroxidation in cultured neurons following exposure to oxygen and glucose deprivation (68, 69). These effects may be result from the sulfhydryl group present in dissolved thiopental. Indeed, we observed an increase in the levels of reduced thiol in the thiopental group (Fig 3A).

## 5 Summary and Outlook

This study was on the differential effects of ketamine and thiopental anesthesia on ultraweak photon emission (UPE) and oxidative-nitrosative stress in rat brains. To our knowledge, no previous study has investigated and compared UPE in the presence and absence of anesthesia. From the obtained results one could conclude that the contrasting effects of ketamine and thiopental on UPE intensity are strongly correlated to the brain’s oxidative stress and antioxidant capacity. These findings may contribute to a broader understanding of how anesthetics influence not only physiological parameters but also the subtle bioenergetic and redox dynamics in neural tissues (70, 71).

Further research is essential to investigate the effects of other anesthetics on the brain. Moreover, exploring the molecular mechanisms underlying the action of the anesthetics, e.g. the role of mitochondria in anesthesia, could provide a deeper understanding of the role of oxidative stress and UPE as biomarkers for anesthetic efficacy and safety. It will provide further insight into how consciousness changes or shuts down during anesthesia, shedding light on the nature of consciousness itself, and help develop safer anesthesia practices, improving patient care and advancing protective strategies for the brain in medical treatments.

## AUTHOR CONTRIBUTIONS

Conceptualization: T.E., V.S., and C.S.; Data curation: N.S. and M.K.G.; Formal analysis: T.E., and V.S., Funding acquisition: T.E., V.S., C.S., D.O.; Methodology: N.S. and M.K.G.; Project administration: T.E. and V.S.; Resources: T.E. and M.K.G.; Supervision: T.E., C.S., and V.S.; Writing an original draft: M.K.G., N.S. revision and editing: V.S., D.O.and C.S.

## References

1. Hemmings HC, Riegelhaupt PM, Kelz MB, Solt K, Eckenhoff RG, Orser BA, Goldstein PA. Towards a comprehensive understanding of anesthetic mechanisms of action: a decade of discovery. Trends in pharmacological sciences. 2019;40(7):464–81.

2. Storm JF, Klink PC, Aru J, Senn W, Goebel R, Pigorini A, et al. An integrative, multiscale view on neural theories of consciousness. Neuron. 2024;112(10):1531–52.

3. Plankar M, Brežan S, Jerman I. The principle of coherence in multi-level brain information processing. Progress in biophysics and molecular biology. 2013;111(1):8–29.

4. Sun Y, Wang C, Dai J. Biophotons as neural communication signals demonstrated by in situ biophoton autography. Photochemical & Photobiological Sciences. 2010;9(3):315–22.

5. Esmaeilpour, T., Fereydouni, E., Dehghani, F. et al. An Experimental Investigation of Ultraweak Photon Emission from Adult Murine Neural Stem Cells. Sci Rep 10, 463 (2020).

6. MacIver BM, Mandema JW, Stanski DR, Bland BH. Thiopental Uncouples Hippocampal and Cortical Synchronized Electroencephalograpbic Activity. The Journal of the American Society of Anesthesiologists. 1996;84(6):1411–24.

7. Arena A, Juel BE, Comolatti R, Thon S, Storm JF. Capacity for consciousness under ketamine anaesthesia is selectively associated with activity in posteromedial cortex in rats. Neuroscience of Consciousness. 2022;2022(1):liac004.

8. Poulet JF, Crochet S. The cortical states of wakefulness. Frontiers in systems neuroscience. 2019;12:64.

9. Salari V, Rodrigues S, Saglamyurek E, Simon C and Oblak D 2022. Are Brain-Computer Interfaces Feasible with Integrated Photonic Chips? Front. Neurosci. 15:780344.

10. Salari V, et al. Relationship between intelligence and spectral characteristics of brain biophoton emission: Correlation does not automatically imply causation. PNAS, 113 (38) E5540–E5541, 2016.

11. Bademosi AT, Steeves J, Karunanithi S, Zalucki OH, Gormal RS, Liu S, et al. Trapping of syntaxin1a in presynaptic nanoclusters by a clinically relevant general anesthetic. Cell reports. 2018;22(2):427–40.

12. Baumgart JP, Zhou Z-Y, Hara M, Cook DC, Hoppa MB, Ryan TA, Hemmings Jr HC. Isoflurane inhibits synaptic vesicle exocytosis through reduced Ca2+ influx, not Ca2+-exocytosis coupling. Proceedings of the National Academy of Sciences. 2015;112(38):11959–64.

13. Bensel BM, Guzik-Lendrum S, Masucci EM, Woll KA, Eckenhoff RG, Gilbert SP. Common general anesthetic propofol impairs kinesin processivity. Proceedings of the National Academy of Sciences. 2017;114(21):E4281–E7.

14. Magistretti PJ, Pellerin L, Rothman DL, Shulman RG. Energy on demand. Science. 1999;283(5401):496–7.

15. Sibson NR, Dhankhar A, Mason GF, Rothman DL, Behar KL, Shulman RG. Stoichiometric coupling of brain glucose metabolism and glutamatergic neuronal activity. Proceedings of the National Academy of Sciences. 1998;95(1):316–21.

16. Moghaddam B, Adams B, Verma A, Daly D. Activation of glutamatergic neurotransmission by ketamine: a novel step in the pathway from NMDA receptor blockade to dopaminergic and cognitive disruptions associated with the prefrontal cortex. Journal of Neuroscience. 1997;17(8):2921–7.

17. McMillan R, Muthukumaraswamy SD. The neurophysiology of ketamine: an integrative review. Reviews in the Neurosciences. 2020;31(5):457–503.

18. Ito T, Suzuki T, Wellman SE, Ho K. Pharmacology of barbiturate tolerance/dependence: GABAA receptors and molecular aspects. Life sciences. 1996;59(3):169–95.

19. Miao N, Nagao K, Lynch C. Thiopental and methohexital depress Ca2+ entry into and glutamate release from cultured neurons. The Journal of the American Society of Anesthesiologists. 1998;88(6):1643–53.

20. Långsjö JW, Salmi E, Kaisti KK, Aalto S, Hinkka S, Aantaa R, et al. Effects of subanesthetic ketamine on regional cerebral glucose metabolism in humans. The Journal of the American Society of Anesthesiologists. 2004;100(5):1065–71.

21. Sokoloff L, Reivich M, Kennedy C, Rosiers MD, Patlak C, Pettigrew K, et al. The [14C] deoxyglucose method for the measurement of local cerebral glucose utilization: theory, procedure, and normal values in the conscious and anesthetized albino rat 1. Journal of neurochemistry. 1977;28(5):897–916.

22. Pospíšil P, Prasad A, Rác M. Role of reactive oxygen species in ultra-weak photon emission in biological systems. Journal of Photochemistry and Photobiology B: Biology. 2014;139:11–23.

23. Pospíšil P, Prasad A, Rác M. Mechanism of the formation of electronically excited species by oxidative metabolic processes: role of reactive oxygen species. Biomolecules. 2019;9(7):258.

24. Dröge W. Free radicals in the physiological control of cell function. Physiological reviews. 2002.

25. Flohé L, Toppo S, Cozza G, Ursini F. A comparison of thiol peroxidase mechanisms. Antioxidants & redox signaling. 2011;15(3):763–80.

26. Ulrich K, Jakob U. The role of thiols in antioxidant systems. Free Radical Biology and Medicine. 2019;140:14–27.

27. Rastogi A, Pospíšil P. Spontaneous ultraweak photon emission imaging of oxidative metabolic processes in human skin: effect of molecular oxygen and antioxidant defense system. Journal of Biomedical Optics. 2011;16(9):096005–7.

28. Kakinuma K, Cadenas E, Boveris A, Chance B. Low level chemiluminescence of intact polymorphonuclear leukocytes. FEBS letters. 1979;102(1):38–42.

29. Murphy K, Birn RM, Bandettini PA. Resting-state fMRI confounds and cleanup. Neuroimage. 2013;80:349–59.

30. Slupe AM, Kirsch JR. Effects of anesthesia on cerebral blood flow, metabolism, and neuroprotection. Journal of Cerebral Blood Flow & Metabolism. 2018;38(12):2192–208.

31. Von Bohlen O, Halbach U. The isolated mammalian brain: an in vivo preparation suitable for pathway tracing. European Journal of Neuroscience. 1999;11(3):1096–100.

32. Gottschalk S, Degtyaruk O, Mc Larney B, Rebling J, Deán-Ben XL, Shoham S, Razansky D. Isolated murine brain model for large-scale optoacoustic calcium imaging. Frontiers in neuroscience. 2019;13:290.

33. Kushikata T, Yoshida H, Kudo M, Salvadori S, Calo G, Hirota K. The effects of neuropeptide S on general anesthesia in rats. Anesthesia & analgesia. 2011;112(4):845–9.

34. Sefati N, Esmaeilpour T, Salari V, Zarifkar A, Dehghani F, Ghaffari MK, et al. Monitoring Alzheimer’s disease via ultraweak photon emission. iScience. 2024;27(1).

35. Ohkawa H, Ohishi N, Yagi K. Assay for lipid peroxides in animal tissues by thiobarbituric acid reaction. Analytical biochemistry. 1979;95(2):351–8.

36. Miranda KM, Espey MG, Wink DA. A rapid, simple spectrophotometric method for simultaneous detection of nitrate and nitrite. Nitric oxide. 2001;5(1):62–71.

37. Levine RL, Garland D, Oliver CN, Amici A, Climent I, Lenz A-G, et al. [49] Determination of carbonyl content in oxidatively modified proteins. Methods in enzymology. 186: Elsevier; 1990. p. 464–78.

38. Rahman I, Kode A, Biswas SK. Assay for quantitative determination of glutathione and glutathione disulfide levels using enzymatic recycling method. Nature protocols. 2006;1(6):3159–65.

39. Kruger NJ. The Bradford method for protein quantitation. The protein protocols handbook. 2009:17–24.

40. Kobayashi M, Takeda M, Sato T, Yamazaki Y, Kaneko K, Ito K-I, et al. In vivo imaging of spontaneous ultraweak photon emission from a rat’s brain correlated with cerebral energy metabolism and oxidative stress. Neuroscience research. 1999;34(2):103–13.

41. Turrens JF. Mitochondrial formation of reactive oxygen species. The Journal of physiology. 2003;552(2):335–44.

42. Tang R, Dai J. Spatiotemporal imaging of glutamate-induced biophotonic activities and transmission in neural circuits. PloS one. 2014;9(1):e85643.

43. Chen Y, Li S, Liang X, Zhang J. Differential alterations to the metabolic connectivity of the cortical and subcortical regions in rat brain during ketamine-induced unconsciousness. Anesthesia & Analgesia. 2022;135(5):1106–14.

44. Hongliang L, Shanglong Y. Effect of thiopental sodium on the release of glutamate and γ-aminobutyric acid from rats prefrontal cortical synaptosomes. Journal of Huazhong University of Science and Technology [Medical Sciences]. 2004;24:602–4.

45. Liu H, Yao S. Thiopental sodium reduces glutamate extracellular levels in rat intact prefrontal cortex. Experimental brain research. 2005;167:666–9.

46. Sonnay S, Duarte JM, Just N, Gruetter R. Energy metabolism in the rat cortex under thiopental anaesthesia measured in vivo by 13C MRS. Journal of neuroscience research. 2017;95(11):2297–306.

47. Chai W, Han Z, Wang Z, Li Z, Xiao F, Sun Y, et al. Biophotonic activity and transmission mediated by mutual actions of neurotransmitters are involved in the origin and altered states of consciousness. Neuroscience bulletin. 2018;34:534–8.

48. Dawson N, Morris BJ, Pratt JA. Subanaesthetic ketamine treatment alters prefrontal cortex connectivity with thalamus and ascending subcortical systems. Schizophrenia bulletin. 2013;39(2):366–77.

49. Del Arco A, Segovia G, Mora F. Blockade of NMDA receptors in the prefrontal cortex increases dopamine and acetylcholine release in the nucleus accumbens and motor activity. Psychopharmacology. 2008;201:325–38.

50. Tao R, Auerbach SB. Anesthetics block morphine-induced increases in serotonin release in rat CNS. Synapse. 1994;18(4):307–14.

51. Tang R, Dai J. Biophoton signal transmission and processing in the brain. Journal of Photochemistry and Photobiology B: Biology. 2014;139:71–5.

52. Lesser J, Koorn R, Vloka J, Kuroda M, Thys D. The interaction of temperature with thiopental and etomidate on extracellular dopamine and glutamate levels in Wistar-Kyoto rats subjected to forebrain ischemia. Acta anaesthesiologica scandinavica. 1999;43(10):989–98.

53. Lambert DG, Willets JM, Atcheson R, Frost C, Smart D, Rowbotham DJ, Smith G. Effects of propofol and thiopentone on potassium-and carbachol-evoked [3H] noradrenaline release and increased [Ca2+] i from SH-SY5Y human neuroblastoma cells. Biochemical pharmacology. 1996;51(12):1613–21.

54. Wang Z, Wang N, Li Z, Xiao F, Dai J. Human high intelligence is involved in spectral redshift of biophotonic activities in the brain. Proceedings of the National Academy of Sciences. 2016;113(31):8753–8.

55. Li Y, Wen G, Ding R, Ren X, Jing C, Liu L, et al. Effects of Single-Dose and Long-Term Ketamine Administration on Tau Phosphorylation–Related Enzymes GSK-3β, CDK5, PP2A, and PP2B in the Mouse Hippocampus. Journal of Molecular Neuroscience. 2020;70:2068–76.

56. Kurian P, Obisesan T, Craddock TJ. Oxidative species-induced excitonic transport in tubulin aromatic networks: Potential implications for neurodegenerative disease. Journal of Photochemistry and Photobiology B: Biology. 2017;175:109–24.

57. Murphy MP. How mitochondria produce reactive oxygen species. Biochemical journal. 2009;417(1):1–13.

58. Venâncio C, Félix L, Almeida V, Coutinho J, Antunes L, Peixoto F, Summavielle T. Acute ketamine impairs mitochondrial function and promotes superoxide dismutase activity in the rat brain. Anesthesia & Analgesia. 2015;120(2):320–8.

59. Chiu C-T, Scheuing L, Liu G, Liao H-M, Linares GR, Lin D, Chuang D-M. The mood stabilizer lithium potentiates the antidepressant-like effects and ameliorates oxidative stress induced by acute ketamine in a mouse model of stress. International Journal of Neuropsychopharmacology. 2015;18(6):pyu102.

60. Schimites P, Segat H, Teixeira L, Martins L, Mangini L, Baccin P, et al. Gallic acid prevents ketamine-induced oxidative damages in brain regions and liver of rats. Neuroscience letters. 2020;714:134560.

61. da Silva FCC, de Oliveira Cito MdC, da Silva MIG, Moura BA, de Aquino Neto MR, Feitosa ML, et al. Behavioral alterations and pro-oxidant effect of a single ketamine administration to mice. Brain research bulletin. 2010;83(1-2):9–15.

62. Scorziello A, Santillo M, Adornetto A, Dell’Aversano C, Sirabella R, Damiano Sa, et al. NO-induced neuroprotection in ischemic preconditioning stimulates mitochondrial Mn-SOD activity and expression via RAS/ERK1/2 pathway. Journal of neurochemistry. 2007;103(4):1472–80.

63. Storz P, Döppler H, Toker A. Protein kinase D mediates mitochondrion-to-nucleus signaling and detoxification from mitochondrial reactive oxygen species. Molecular and cellular biology. 2005;25(19):8520–30.

64. Vogel C, Silva GM, Marcotte EM. Protein expression regulation under oxidative stress. Molecular & Cellular Proteomics. 2011;10(12).

65. Buttgereit F, Brand MD. A hierarchy of ATP-consuming processes in mammalian cells. Biochemical Journal. 1995;312(1):163–7.

66. Schwer CI, Lehane C, Guelzow T, Zenker S, Strosing KM, Spassov S, et al. Thiopental inhibits global protein synthesis by repression of eukaryotic elongation factor 2 and protects from hypoxic neuronal cell death. PloS one. 2013;8(10):e77258.

67. Chatterjee S. Oxidative stress, inflammation, and disease. Oxidative stress and biomaterials: Elsevier; 2016. p. 35–58.

68. Inoue G, Ohtaki Y, Satoh K, Odanaka Y, Katoh A, Suzuki K, et al. Sedation Therapy in Intensive Care Units: Harnessing the Power of Antioxidants to Combat Oxidative Stress. Biomedicines. 2023;11(8):2129.

69. Almaas R, Saugstad OD, Pleasure D, Rootwelt T. Effect of barbiturates on hydroxyl radicals, lipid peroxidation, and hypoxic cell death in human NT2-N neurons. The Journal of the American Society of Anesthesiologists. 2000;92(3):764–74.

70. Salari V et al. Ultraweak Photon Emission in the Brain. J. Integrative Neuroscience. 14, 3, 419–429 (2015).

71. Esmaeilpour T, et al. Effect of methamphetamine on ultraweak photon emission and level of reactive oxygen species in male rat brain. Neuroscience Letters, 801, 137136, 2023.

